# Breaking the species barrier: recurrent genomic introgressions from very distant lineages in a ciliate

**DOI:** 10.64898/2026.02.20.707023

**Authors:** Florian Bénitière, Olivier Arnaiz, Simon Penel, Xue Zhang, Sandra Duharcourt, Eric Meyer, Linda Sperling, Laurent Duret

## Abstract

The process of speciation, through which two reproductively isolated species emerge from an ancestral population, is often gradual, including a period during which gene flow still occurs, and hence speciation remains reversible. However, if isolation persists, the progressive accumulation of genetic incompatibilities ultimately locks the speciation process, reaching the point where hybrid offspring are inviable or sterile. In plants and animals, the establishment of species barriers is quite rapid: beyond a net genetic divergence of 0.02 substitutions per synonymous site, interspecific introgressions become extremely rare. Here we report that in ciliates, species barriers can be broken despite huge genetic divergence. We show that the genome of *Paramecium sonneborni* contains hundreds of DNA segments that were acquired by recurrent horizontal transfers from multiple very distant lineages (synonymous divergence ∼1 substitution/site). These segments approach the size of full-length chromosomes, which indicates that they were introgressed by inter-specific mating. These data show that in the ancestry of *P. sonneborni*, hybridization events with highly divergent species occurred repeatedly and led to viable and fertile offspring. We propose that this unexpected promiscuous sexual behavior is made possible by the fact that paramecia separate somatic and germline functions in two types of nuclei: the elimination of non-self DNA fragments during the development of the somatic genome allows individuals of a species to remain phenotypically the same, despite recurrent genetic flow from distant species into their germline.

**Significance Statement:** An important issue in evolutionary biology is to understand the process of speciation, i.e. how two reproductively isolated species emerge from an ancestral population. Various factors can initiate the speciation process, but whatever the initial factor, once two populations become reproductively isolated, they start accumulating genetic incompatibilities. In plants and animals, these incompatibilities generally accumulate rapidly, preventing the formation of viable and fertile hybrids, except between genetically closely related species. We show here that this rule does not apply to all eukaryotes. We discovered that the unicellular eukaryote *Paramecium sonneborni* recurrently acquired large DNA segments via hybridization with very distantly related species. This unexpected phenomenon highlights the diversity of factors that affect the delineation of species across eukaryotes.

## Introduction

An important issue in evolutionary biology is to understand the process of speciation, i.e. how two reproductively isolated species emerge from an ancestral population. Various factors can initiate the speciation process (e.g. isolation by geographic barriers, adaptation to distinct ecological niches, etc.), but whatever the initial factor, once two populations become reproductively isolated from each other, they start evolving independently, and hence progressively accumulate genetic incompatibilities. These so-called Bateson–Dobzhansky–Muller incompatibilities (BDMIs) decrease the fitness of hybrid individuals, and ultimately lead to their sterility or non-viability (1). The speciation process is often gradual, i.e. includes a ‘grey zone’ during which genetic introgressions are still possible between “semi-isolated” species (2). Thus, if secondary contacts occur during the early phase, then hybridizations may restore genetic homogeneity, thereby stopping the speciation process. However, as BDMIs accrue, selection is expected to favor the evolution of traits that prevent interspecific mating, thereby further reinforcing genetic isolation (3). The rate at which reproductive barriers accumulate varies widely across taxa (4, 5). Among the multiple factors that drive this variation, the parental genetic distance appears to be a key parameter (5, 6). In particular, the analysis of population genomics data from a wide range of plants and animals has shown that introgressions are restricted to species that are genetically very close: in animals, gene flow appears to be abolished as soon as the net synonymous divergence exceeds 0.02 substitutions per site, and this threshold is even lower in plants (7, 8). Thus, although hybridization is common in nature, it generally leads to inviable or sterile offspring, and hence does not contribute to gene flow, except between genetically closely related species (9).

Here we report an extreme exception to this rule, that we discovered incidentally in the course of sequencing the genomes of several *Paramecium* species (10). We present evidence of recurrent events of introgression within the genome of *Paramecium sonneborni*, originating from very remote species (synonymous divergence ∼1 substitution/site), and we propose a model explaining how fertile hybrids can be formed between such distant species, despite BDMIs.

To facilitate the interpretation of our observations, it is useful to give some background information on the biology and genetics of *Paramecium* species. These are aquatic unicellular eukaryotes that are commonly found in ponds, where they feed on bacteria, yeast, or algae. They were among the first ciliates to be observed, thanks to the invention of the microscope in the late 17^th^ century, and since then have become a widely used model for biological studies (11). Their life cycle includes a vegetative phase (reproduction by mitotic cell divisions), followed by a sexual phase, which involves either conjugation (the pairing and reciprocal fertilization of two cells of complementary mating types) or autogamy (a self-fertilization process, present in all species of the *P. aurelia* clade) (12).

Like all ciliates, *Paramecium* cells possess two types of nuclei: a highly polyploid somatic macronucleus (MAC), responsible for gene expression, and a diploid germline micronucleus (MIC), which is not expressed, but where meiosis occurs. At each sexual generation, the parental MAC is lost, while new MIC and MAC develop from copies of the zygotic nucleus. During MAC development, many DNA segments are reproducibly eliminated, thus leading to a somatic genome that is smaller than the germline genome (13, 14).

The genus *Paramecium* encompasses a wide diversity of species. Based on cytological observations, 19 species can be distinguished (15). However, genetic crosses have revealed a large amount of cryptic diversity within these groups. Notably, early studies have established that *P. aurelia*, initially considered as one ‘species’, was in fact a complex of 14 sibling species (‘syngens’) that are morphologically indistinguishable but sexually incompatible (16). Currently, 20 species have been ascribed to the *P. aurelia* species complex (17): 15 that are morphologically indistinguishable (named *P. primaurelia* to *P. quindecaurelia*), together with *P. sonneborni*, *P. schewiakoffi*, and three sibling species of *P. jenningsi* (16, 18–21). Some of these species show varying degrees of sexual incompatibility – being still able to form conjugant pairs, and the analysis of polymorphism data on a few genes revealed some incongruities, suggesting that hybridizations might occur (22, 23). Nevertheless, phylogenetic and population genetic studies based on MAC genome sequences have confirmed that these syngens correspond to *bona fide* species, reproductively isolated and genetically divergent (for a review, see (24)). The phylogeny of this clade is now well resolved, with robust and congruent trees obtained both from nuclear and mitochondrial genes (17, 25, 26). This species complex encompasses organisms that are genetically as distant as the most remotely related mammals (17, 27).

Little is known regarding the ecological factors that may have driven the diversification of this clade. Some species are preferentially found in tropical environments (e.g. *P. sexaurelia, P. octaurelia, P. tredecaurelia, P. quadecaurelia,* and *P. sonneborni*), or in temperate or cold climate zones (e.g. *P. biaurelia* and *P. novaurelia*), whereas others are totally cosmopolitan (e.g. *P. primaurelia* or *P. tetraurelia*) (28). Overall, most species are present on several continents and overlap in their geographical distribution (28). The sequencing of 139 strains of widespread geographical origin showed that the genome-wide level of neutral polymorphism is quite low within each *aurelia* species (ranging from 0.6% in *P. tetraurelia* to 2.3% in *P. sexaurelia*), probably because of a high rate of autogamy (29, 30). One noteworthy observation is that three rounds of whole genome duplications (WGDs) occurred in the ancestry of the *aurelia* lineage, and the last one shortly predates the radiation of the complex (25, 31, 32). The differential loss of paralogous gene copies after a WGD is expected to be an important source of BDMIs (31, 33), and hence it has been suggested that these WGDs might have contributed to the diversification of the *P. aurelia* complex (25, 31, 32). The process of somatic DNA rearrangements is also suspected to cause BDMIs in ciliates (34).

To study the organization and evolution of the germline genome, we sequenced the MIC of 7 *P. aurelia* species (27, 10). We obtained a high-quality assembly for the *P. tetraurelia* MIC genome (104 Mb), which consists of ∼160 tiny chromosomes (300 kb – 1.2 Mb) (10). Three genomic compartments can be distinguished, depending on their fate during the development of the new polyploid MAC: i) MIC-limited regions, that are eliminated from the MAC, ii) constitutive MAC-destined regions, that are systematically retained in the MAC, and iii) MAC-variable regions, which correspond to DNA sequences that are retained in only a small fraction of MAC copies (27). These three compartments represent respectively 20%, 71% and 9% of the MIC genome. The MAC-destined regions are very gene-rich (∼500 genes per Mb), whereas most repeated sequences are located in the MIC-limited regions (27, 10). Two types of MIC-limited regions can be distinguished: the ‘Internal Eliminated Sequences’ (IESs, totalizing 3.9 Mb) that correspond to short sequences (typically <150bp) interspersed within MAC-destined regions and notably within genes, whereas the other eliminated sequences (16 Mb) are generally located towards the end of MIC chromosomes (10). Most other species show similar genome content, with a MIC genome ∼100 Mb. The only exception is *P. sonneborni*, where the MIC reaches 217 Mb (10). The size of the MAC-destined compartment is similar across species, ranging from 65 Mb in *P. tredecaurelia* to 82 Mb in *P. sonneborni* (27). Thus, the large size of the *P. sonneborni* MIC results from a huge inflation of its MIC-specific compartment.

To investigate the cause of this increase, we compared *P. sonneborni* MIC sequences to other *Paramecium* genomes, and discovered that *P. sonneborni* contained long regions with very high sequence similarity to distantly related *aurelia* species. This prompted us to systematically investigate patterns of horizontal transfer across paramecia. We found evidence of recurrent events of introgression of chromosome-size segments within the MIC-specific genomic compartment of *P. sonneborni*, originating from very distant species. We propose that these introgressions result from allotetraploidization events, and we discuss the mechanism that could explain how these events can be tolerated despite the accumulation of BDMIs.

## Results

### Evidence of massive horizontal transfer in the exceptionally large MIC genome of P. sonneborni

To examine the origin of the large MIC-limited compartment in *P. sonneborni*, we compared its sequences to all available genomes from 13 species of the *P. aurelia* complex (7 species for which both MIC and MAC genomes have been sequenced, and 6 species for which only the MAC is available). The species phylogeny, inferred from protein-alignments of 473 single-copy orthologs (17), indicates that *P. sonneborni* is closely related to *P. jenningsi*, but quite distant from all the other species that we analyzed (Figure 1A). To estimate the neutral divergence between *P. sonneborni* and other *aurelia* species, we measured the substitution rate at synonymous sites (dS). The mean dS is 0.27 substitution per site for *P. jenningsi*,, whereas it reaches 0.8 to 1.0 for the other species (Suppl. Fig. S1). This implies that outside of coding regions, sequences are generally too divergent to be aligned across these species. Yet, we noticed that several *P. sonneborni* MIC contigs contain long segments with very high similarity to distantly related species. For instance, we observed a *P. sonneborni* contig of 100kb presenting long stretches above 99% identity with a MAC scaffold from *P. quadecaurelia*, a level of similarity higher than with any other *aurelia* species, including *P. jenningsi* (Figure 1B). This suggests that this long genomic segment has been subjected to horizontal transfer (HT) between the *P. sonneborni* and *P. quadecaurelia* lineages.

**Figure 1.**
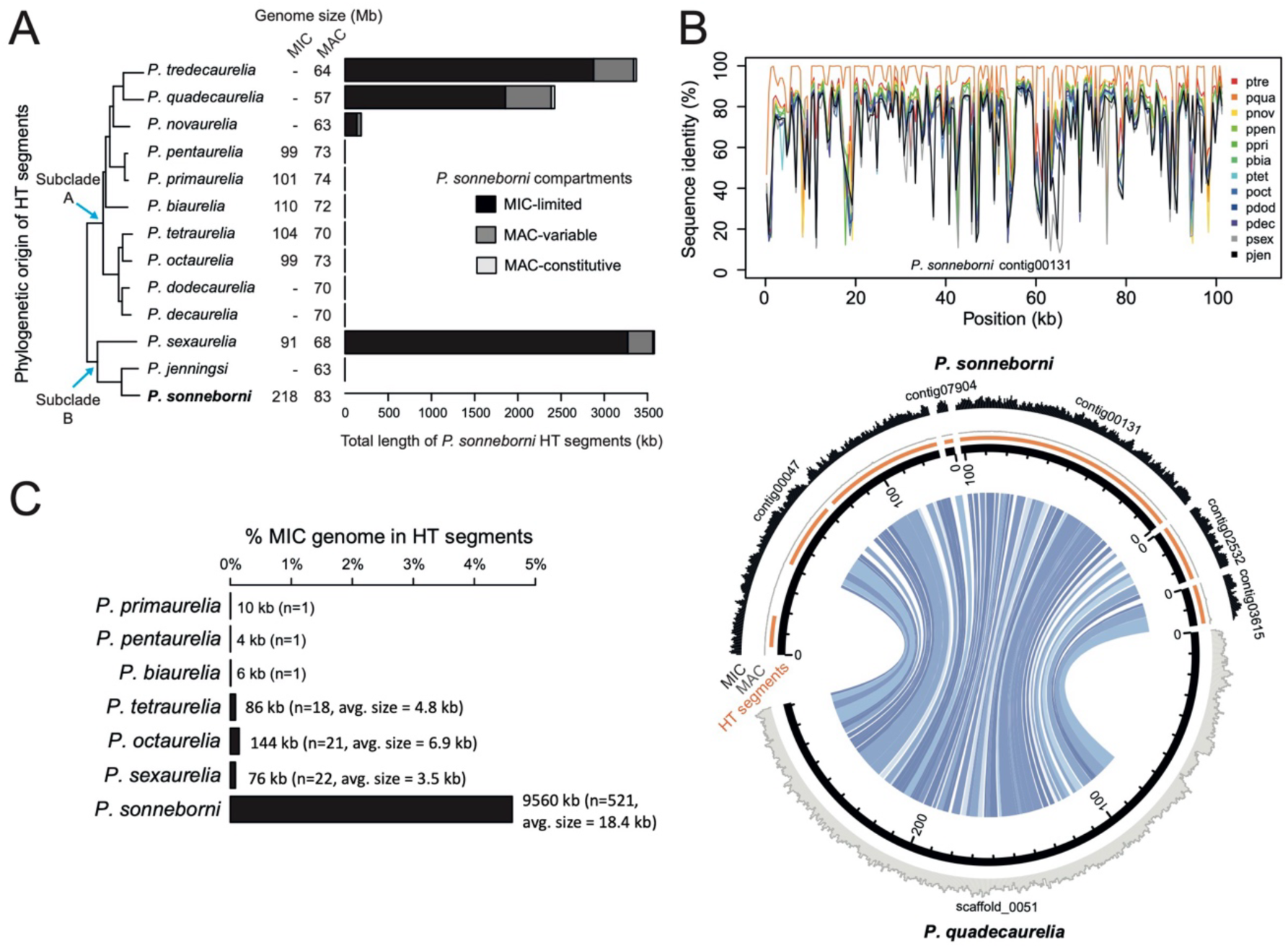
Detection of horizontally transferred DNA segments in Paramecia. **A.** Phylogenetic tree of *Paramecium aurelia* species (modified from (25)). This tree was inferred from protein alignments of 473 single-copy orthologs, and all nodes show very strong bootstrap support (25). For each species, the sizes of the MAC and, if available, MIC genome assemblies are indicated. All the *aurelia* species have MAC genomes around 70 Mb and MIC genomes around 100 Mb in size, with the notable exception of *P. sonneborni* whose MIC genome is more than twice as large (10). The barplot indicates the phylogenetic origin of HT segments identified in the *P. sonneborni* genome. HT regions can be localized in three genomic compartments: MIC-limited (black), MAC-variable (dark grey) and MAC-destined (light grey). **B.** Example of a region of the *P. sonneborni* MIC genome that presents the hallmarks of HT from other *P. aurelia* species (figures for all other HT segments are available in the Zenodo repository). The upper panel presents the data for *P. sonneborni* contig00131: the contig was cut into 500bp-long non-overlapping windows, and each window was compared (BLASTN) to all other *P. aurelia* MAC genomes. Each colored line indicates the level of sequence identity with their best hit in the corresponding target genome (in the color legend, species are listed in the same order as in the tree from panel A). HT segments were identified in *P. sonneborni* by searching for sets of neighboring windows with conserved synteny, and whose best hits were not with its sister group (*P. jenningsi*, pjen) (see Methods for details). In this example, the entire *P. sonneborni* contig shows a very high level of similarity with the genome of *P. tredecaurelia* (ptre), more than with any other *P. aurelia* species. Four digit species codes refer to the first letters of the genus and species names (e.g. pson for *P. sonneborni*). Lower panel, the circos plot shows the conservation of synteny between five *P. sonneborni* contigs containing HT segments (orange) and the putative donor region of the *P. quadecaurelia* MAC genome (scaffold_0051). Histograms describe DNA-seq coverage for MIC DNA sample (black) and MAC DNA sample (grey). In this example, the HT segments are located in the MIC-limited compartment of *P. sonneborni*: they are well covered in the MIC DNA sample but not in the MAC. The colors of the arcs are defined by the similarity between regions. **C.** Percentage of MIC genomes in horizontally transferred (HT) segments. The numbers next to each bar indicate the cumulative size and number of HT segments.

This unexpected observation led us to systematically analyze all MIC genomes (from 7 species) to search for loci presenting evidence of recent HT within the *aurelia* complex. For this, we divided each MIC genome (query) into non-overlapping 500 nt windows and compared (with blastn) each window with all other available *Paramecium* genomes (targets) (Figure 1A). We then searched for sets of neighboring windows (at least 2 kb-long in total) with conserved order and orientation between the query and the target, and showing a strong signal of HT (i.e. a level of sequence similarity significantly higher with a genome that is not the sister group of the focal species, than with any other *aurelia* species).

In most species we identified only a limited number of HT segments (1 to 22 segments, totalizing up to 144 kb; Figure 1C). Strikingly, in the *P. sonneborni* MIC genome, we identified 521 HT segments totaling almost 10 Mb. HT segments are on average 2 to 5 times longer in *P. sonneborni* (mean length = 18.4 kb) than in other species (Figure 1C). In *P. sonneborni* we detected 120 HT segments longer than 25 kb (totaling 5 Mb). The length of HT segments in *P. sonneborni* is probably underestimated because the MIC assembly is quite fragmented (N50 ∼33 kb, (10)). Indeed, we found several cases where a single region in the MAC genome of a target species showed synteny with multiple *P. sonneborni* contigs (Figure 1B). These observations indicate that many HT segments in *P. sonneborni* are very long, approaching the size of an entire chromosome (>220 kb for the example shown in Figure 1B). Conversely, none of the HT segments in the other species were larger than 25 kb.

Several observations allowed us to exclude the hypothesis of a DNA contamination artefact. First, contamination with DNA from the target genomes should lead to nearly identical sequences, whereas for many HT segments, the level of sequence divergence ranges between 1% and 12% (Figure 2), far too high to result from sequencing errors. Second, two of the target genomes with which we detected many HT segments (*P. quadecaurelia* and *P. novaurelia;* Figure 1A) were sequenced in the USA (25), and were never handled in the laboratory where *P. sonneborni* MAC and MIC DNA were sequenced (Genoscope, France). Finally, the sequencing depth along *P. sonneborni* HT segments is the same as that of the rest of the genome, which would not be expected if they resulted from contaminant DNA (Suppl. Fig. S2).

**Figure 2.**
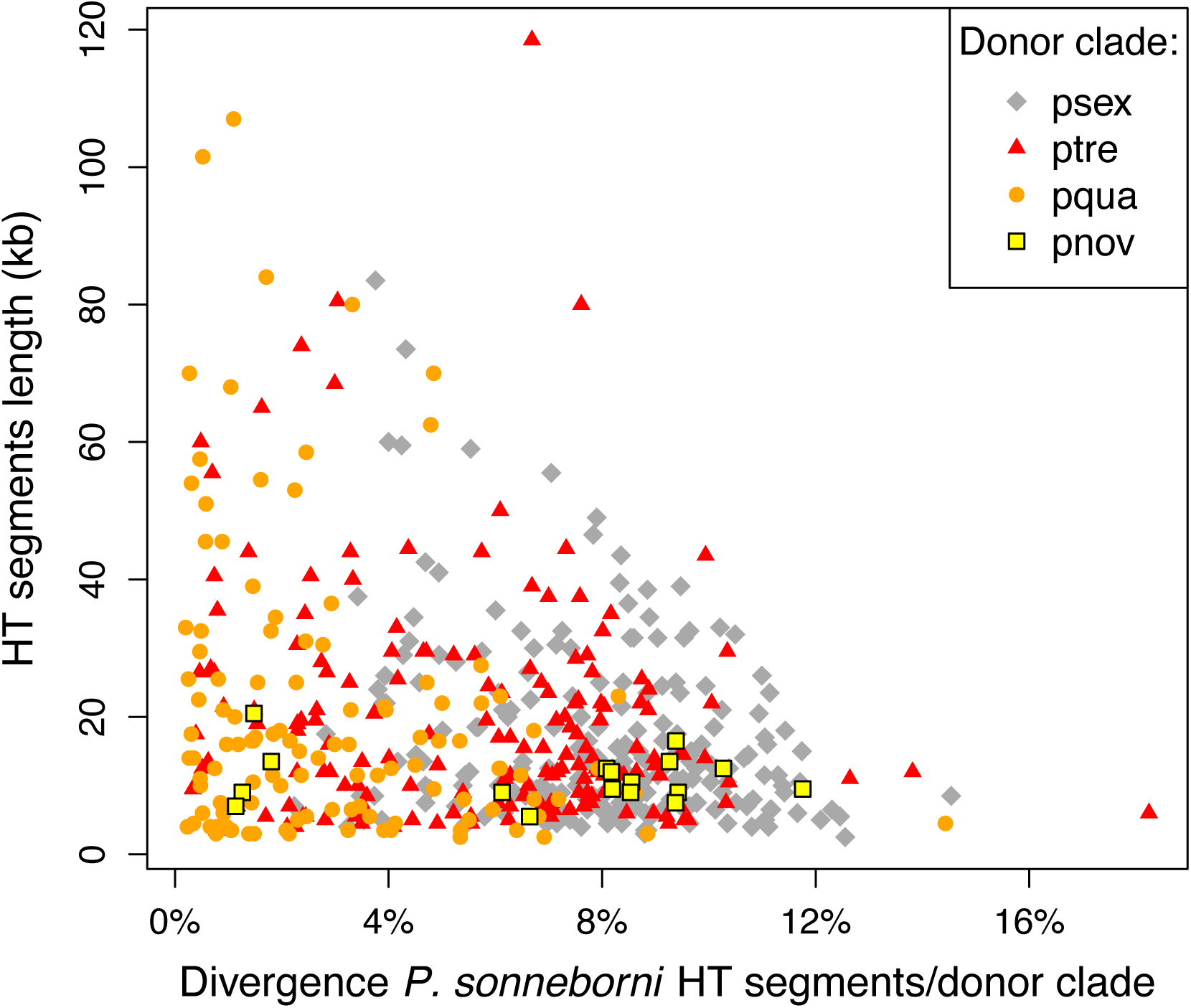
Relationship between the size of HT segments in the *P. sonneborni* genome, and their divergence from the donor lineage. The origin of the donor clade is indicated by different colors and dot styles. Species codes as in Figure 1.

The HT segments detected in *P. sonneborni* involve four distinct lineages (represented by *P. sexaurelia, P. tredecaurelia, P. quadecaurelia,* and *P. novaurelia*) (Figure 1A). Among HT segments detected by comparison between *P. sonneborni* and *P. sexaurelia* (the only species pair for which we had both MIC genome assemblies), we observed many conserved IESs (Suppl. Fig. S3), which demonstrates that these HT events involved the transfer of MIC DNA and not of MAC DNA (where IESs are absent). *P. sexaurelia* belongs to the same subclade as *P. sonneborni* but is nevertheless very distant (average dS=0.8; (27)). The three other lineages belong to subclade A, very divergent from *P. sonneborni* (dS∼1; (27)). We did not detect any HT segment with the other lineages of subclade A (Figure 1A).

The MIC and MAC reference genome assemblies of *P. sonneborni* derive from the reference strain (ATCC 30995), which was isolated in Texas in the early 1980s (18). To assess whether the HT events we detected might result from some weird accident during the lab history of this specific strain, we sequenced another strain (Rio Lg_Jac 5III, hereafter termed ‘Rio Lg’), which was collected in Brazil in 2012. These two strains are genetically close (1.2 SNPs per kb). We mapped Rio Lg reads on the reference MIC genome assembly (ATCC 30995). Given that we sequenced whole-cell DNA extracts and that the ploidy is ∼400 fold higher in the MAC than in MIC nuclei, the sequencing depth of MIC-specific regions was very limited: only 6.7% of this compartment is covered by at least one read. Nevertheless, out of 521 HT segments, 65 (12.5%) were retrieved in Rio Lg. As expected, the fraction of HT segments that were retrieved in different genomics compartments reflects the corresponding sequencing coverage in Rio Lg (Suppl. Table S1), and the different lineages involved in these HT events were in similar proportion as in ATCC 30995 (Suppl. Table S2). We therefore conclude that the HT events detected in the MIC genome of *P. sonneborni* are not an oddity of the ATCC 30995 reference strain, but are also present in other strains of geographically distant origin.

It should be noted that MIC genome sequences are available for only 7 of the 13 analyzed species. Because of this sampling limitation, HT segments present in the MIC genome of the query were detectable only if they encompassed MAC-destined regions in the target genome (see Methods for details). Hence, the set of segments that we identified is probably an underestimate of the total amount of HT.

### P. sonneborni is the recipient of recurrent horizontal transfers from distantly related lineages

The above results show that gene flow occurred between *P. sonneborni* and several other *Paramecium* lineages. To establish the direction of these transfers, we analyzed the distribution of HT segments in the MIC of *P. sonneborni*: 85% are located in the MIC-limited compartment of *P. sonneborni*, 14% in MAC-variable regions and only 1% in MAC-destined regions (these compartments represent respectively 50%, 14% and 36% of the MIC genome) (Figure 1A). Thus, the majority of HT segments are confined to genomic compartments that are either eliminated or variably retained during MAC development. As a result, although the HT segments correspond to MAC-destined regions in the target genomes and are therefore very gene-rich, the genes they contain are unlikely to be expressed in *P. sonneborni*. In total, among the 521 HT segments, we identified only 35 genes located in the MAC-destined compartment of *P. sonneborni* (Suppl. Table S3). Comparison with their homologs in other *aurelia* species showed that at least 25 of them are pseudogenes (containing nonsense or frameshift mutations), and among the ten that appear to be intact, only four show evidence of transcription (> 1RPKM) in vegetative cells. Thus, even though HT segments detected in *P. sonneborni* included several thousand genes, most of them have lost their functionality. This implies that *P. sonneborni* was the recipient, and not the donor of the horizontal transfers.

We did not detect any case of a gene from the donor genome integrated by recombination at its homologous locus in the *P. sonneborni* MIC genome. Most of the HT segments are located in the MIC-limited compartment, and hence away from genic regions, and the few (pseudo)genes that we detected in the MAC-destined compartment are most often located towards the end of MAC chromosomes (Suppl. Table S3). Thus, these HT events did not lead to the replacement of recipient genes by their donor homologues, but to the addition of new gene copies (i.e. gene duplication via HT).

*P. sonneborni* has received HT from four distinct donor lineages (Figure 1A). Several observations indicate that the detected HT segments result from numerous independent genetic transfers, and not from just four occasional events. Indeed, the level of sequence divergence observed between *P. sonneborni* HT segments and the target genomes varies widely among HT segments, ranging from ∼0% to ∼12%, even among segments coming from the same lineage (Figure 2). This implies that multiple HT events occurred from each lineage. Two non-exclusive hypotheses can be proposed to explain this large variance: i) some HT events are more ancient than others, and hence have had more time to accumulate divergence between the donor and the recipient lineages, or ii) HT segments have about the same age, but originated from multiple related species that vary in their level of divergence from the reference species used in our analyses. One noteworthy observation is that the length of HT segments tends to decrease as the sequence divergence increases (Figure 2): for a given donor lineage, the longest HT segments (approaching the size of full-length chromosomes) are generally less divergent from the reference target than other HT segments, whereas the most divergent HT segments are always relatively short. The most parsimonious explanation for this pattern is that after the transfer of full-length chromosomes, HT segments progressively shorten via chromosomal rearrangements (inversions, translocations, deletions) and concomitantly diverge via point substitutions. This therefore suggests that the variance in level of divergence mainly reflects the variability in the age of HT events. In the absence of a fossil record, it is difficult to calibrate a molecular clock to date HT events. However, we can use the level of sequence divergence as a ‘relative time unit’, i.e. to estimate the frequency at which HT events occur, relative to nucleotide substitutions. If we consider the most recent HT segments from each donor lineage, their levels of divergence are respectively 0.2% for of *P. quadecaurelia*, 0.3% for *P. tredecaurelia*, 1.1% for *P. novaurelia* and 2.2% for *P. sexaurelia* (Figure 2). This indicates that during a time period corresponding to ∼2% of sequence divergence, at least four distinct events occurred in the *P. sonneborni* ancestry. By extrapolation, this suggests that the HT segments that we detected, with divergence levels reaching 12% or more, resulted from tens of independent events.

Little is known regarding the ecology of *P. sonneborni*. Although this species is rare, it is distributed across very wide geographical areas (the four strains reported so far are from Texas, Dominican republic, Cyprus and Brazil) (18, 35, 36). The high frequency of introgression coming from *P. sexaurelia, P. tredecaurelia*, and *P. quadecaurelia* lineages might be due to the fact that like *P. sonneborni*, these species are preferentially found in warm climate zones (28).

In principle, when two species are able to form hybrids, introgressions may occur in both directions. Unfortunately, we could not assess the presence of HT segments related to *P. sonneborni* in any of the four lineages from which it received DNA (represented in our data set by *P. tredecaurelia, P. quadecaurelia, P. novaurelia* and *P. sexaurelia*): for the first three, we do not have a MIC genome assembly, and for *P. sexaurelia*, we do not have an appropriate sister group to detect HT from *P. sonneborni*. Thus, the question of whether transfers occur in both directions remains to be investigated.

## Discussion

### Recurrent hybridization with genetically highly divergent species

Different mechanisms can mediate the horizontal transfer (HT) of genetic material from one species to another. In eukaryotes, a distinction is made between ‘introgressions’, i.e. HTs that occur via sexual processes (involving mating, fusion of gametes and meiotic recombination) and other forms of HTs, that do not involve inter-specific mating (e.g. via viral vectors, or by direct uptake of DNA from the environment – including from endosymbionts) (37–39). It is well established that non-sexual HT events can occur between very distant taxa, spread across the tree of life (e.g. some insects received HTs from bacteria, plants or fungi (40, 41)). But introgressions can occur only between species that are close enough to mate and produce fertile hybrids. Although hybridizations are quite common in nature, they often lead to inviable or sterile offspring (9). Different types of processes cause this decreased fitness of hybrids. First, after their divergence, sister species tend to accumulate genetic incompatibilities (i.e. alleles that are neutral or beneficial in one species but that interact negatively with the alleles in the other species), so-called Bateson–Dobzhansky– Muller incompatibilities (BDMIs) (1). Notably, as each species has adapted to its specific ecological niche, the mixed set of alleles in the hybrids may turn out to be unfit in both environments. Second, chromosomal rearrangements and sequence divergence impair homologous recombination, and thereby cause incorrect chromosome segregation during meiosis in hybrids (42). Hence, the probability of forming fertile hybrids (and the potential of inter-species gene flow) decreases with increasing genetic distance (9). Typically, in animals and plants, the amount of gene flow across populations/species drops drastically beyond 2% of net synonymous divergence (7, 8) and in budding yeast, hybrid viability falls to almost zero above 10% of genetic divergence (42).

Here we analyzed MIC genomes from 7 species of the *P. aurelia* complex to search for loci presenting evidence of recent HT. In most species, we detected very few loci for which the observed genetic distances are inconsistent with the species phylogeny, as expected for reproductively isolated species. But we show that in *P. sonneborni*, at least 5% of the genome has been acquired via recurrent events of horizontal transfer from multiple distant lineages. Two lines of evidence indicate that these transfers occurred via inter-specific hybridization. First, the size of the longest HT segments that we detected in *P. sonneborni* (> 200 kb) is in the range of germline chromosome sizes (300 kb to 1 Mb) (10), which implies that they were most probably transferred as full-length chromosomes. Second, HT segments encompass IESs, which indicates that the transferred DNA derived from the MIC. Thus, the most likely explanation is that these HT events result from inter-specific mating and conjugation. Alternative hypotheses involving transfers mediated by viruses or other mobile elements are not compatible with the observed length of HT segments, since the carrying capacity of viral or viral-like particles is generally much lower (43). In protists that feed on other organisms via phagocytosis, like Paramecium, it has been proposed that HT might simply be a by-product of the rupture of digestive vacuoles, releasing DNA from their prey (37, 44). Little is known about the ecology of *P. sonneborni*, but even if that species was able to feed on other paramecia, given that the ploidy is ∼400 fold higher in the MAC than in MIC nuclei, this putative source of transfer would be expected to consist essentially of MAC DNA, in contradiction with our observation of a MIC origin. Our results therefore imply that the *P. sonneborni* lineage recurrently mated and produced fertile offspring with very distant species (synonymous divergence: dS∼0.8 to 1 substitution per site). For comparison, this corresponds to the genetic distance between the most divergent mammalian taxa (e.g. monotremes vs. placental mammals (45)).

Known examples of distant species able to produce viable hybrid offspring are quite rare. The three most extreme cases reported so far are two species of fern that diverged ∼60 Mya (46), sturgeon and paddlefish that diverged ∼184 Mya (47), and two gar fish species that diverged ∼100 Mya (48). The latter case is particularly striking, because unlike the other two, the hybrids are not only viable, but also fertile (48). It has been suggested that the absence of strong BDMI in gars results from the fact that their genomes evolve extremely slowly (48). The same explanation probably holds for the two other cases: the median dS between sturgeon and paddlefish is only 0.12 (49), and in the fern, the dS between the two parental genomes is 0.1 (46). Thus, even though these species have been separated for a very long period, they are genetically quite close. Although introgressions are rare at this level of genetic divergence (7, 8), several cases have been reported. For instance, gene flow has been observed in the *Drosophila yakuba* clade (∼3 Mya of separation), or in *Ciona* (sea squirts; 1.5-2 Mya of separation), between species showing a dS of 0.12 to 0.14 (50, 51). To our knowledge, *P. sonneborni* is the first reported case of a eukaryote showing promiscuous sexual behavior with genetically highly diverged species (dS∼1).

### Remaining the same despite hybridization with distant species

Our observations raise three main questions. First, it is unclear how such distant species could have successfully interbred. In standard lab conditions, *P. sonneborni* was not observed to conjugate with any other *P. aurelia* species, not even with the closely related *P. jenningsi* (18, 35). More extensive analyses in other species concluded that inter-specific conjugation may be blocked at a number of different steps: mating type recognition and agglutination, formation of conjugating pairs, cross-fertilization, and development of functional zygotic MACs. Although all *P. aurelia* species have two homologous mating types (O and E), inter-specific agglutination of O and E cells is only observed in some pairs of species from the same subclade, and in most cases it does not proceed to the formation of conjugating pairs (16). Inter-specific transformation experiments have shown that the E-specific transmembrane protein mtA is directly responsible for the specific recognition of O cells of the same species (52). In the lab, the need for mating type recognition can be bypassed by chemical treatments that induce the successful conjugation of a single mating type with itself (53). Chemical treatments can also induce the formation of conjugating pairs between species (e.g. between a *P. aurelia* species and the outgroup *P. multimicronucleatum*), but in such cases genetic exchange appears to fail and surviving ex-conjugants likely result from self-fertilization (53). Even in the best of cases where inter-specific conjugation can occur naturally, e.g. between the closely related *P. tetraurelia* and *P. octaurelia*, most ex-conjugants do not survive, genetic exchange rarely succeeds, and the rare viable true hybrids are all completely sterile (54).

While the origin of the *P. sonneborni* hybrid ancestors thus remains mysterious, a second question is how they could escape the negative consequences of BDMIs. Here the answer may lie in the peculiar nuclear dimorphism of ciliates. Indeed, the HT segments identified in the genome of *P. sonneborni* are essentially confined to its MIC-limited compartment and hence are not expressed, which effectively hides genetic incompatibilities. In *P. aurelia* species, the distinction between segments that are retained or eliminated during MAC development relies on a comparison between the maternal MIC and the maternal MAC, via small RNAs that are produced from the entire MIC genome during meiosis and are still present in the cytoplasm of the zygote after conjugation: the so-called scnRNAs mediate a genomic subtraction that targets the elimination from the new developing MAC of all maternal MIC sequences that are absent from the maternal MAC (13, 14) (Figure 3). We conjecture that in *P. sonneborni*, a similar mechanism also leads to the elimination of sequences that are present in the paternal MIC but absent from the maternal MAC. This would explain how, after conjugation of *P. sonneborni* with a distantly related species, the chromosomes from the latter may be systematically eliminated from the developing MAC in the F1 hybrid carrying the *P. sonneborni* cytoplasm. That the current species indeed derives from that side of the cross after each inter-specific mating is indicated by the *P. sonneborni* mitochondrial genome, whose phylogenetic position is in total agreement with the species tree inferred from MAC nuclear genomes (17, 26) (Figure 3). In other words, the successful matings of *P. sonneborni* ancestors with distant relatives may have been facilitated by the capacity of *P. aurelia* species to exclude non-self DNA from the zygotic MAC (self DNA being defined by the maternal MAC genome). Interestingly, a similar situation has been reported in a dipteran insect, which possesses two chromosomes that have been acquired by introgression from a distant lineage, but that are restricted to the germline, and hence are not expressed in the soma (55).

**Figure 3.**
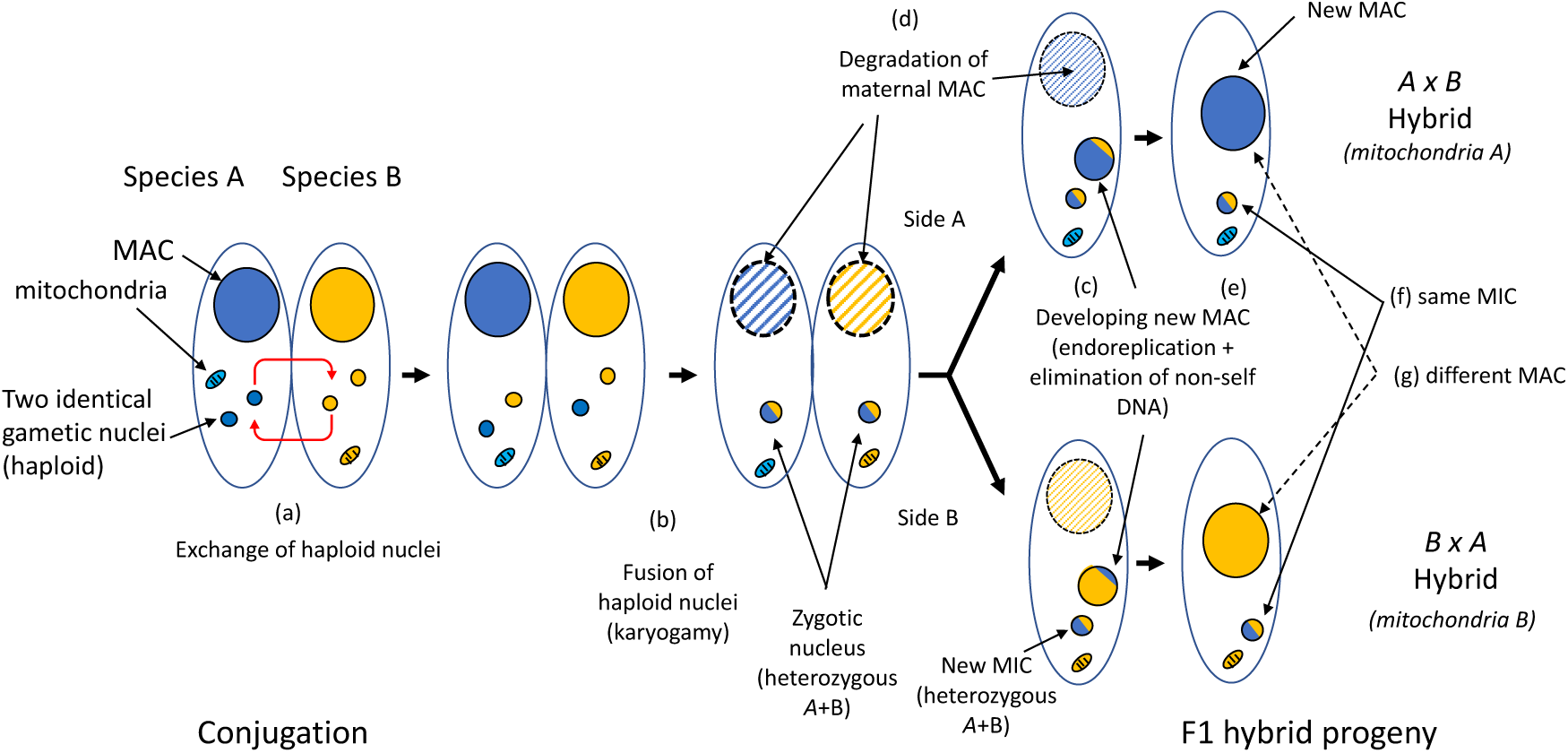
Model: how the nuclear dimorphism in Paramecia might hide interspecific genetic incompatibilities. Sexuality in paramecia occurs through conjugation: cells with compatible mating types pair and engage meiosis. Three of the four resulting haploid nuclei are eliminated, and the remaining one undergoes a mitotic division, leading to the formation of two identical haploid gametic nuclei. The two paired cells exchange one haploid nucleus (a), and then, in each cell, the haploid nuclei fuse to form the new zygotic nucleus (b). After two mitotic divisions, one nucleus differentiates to form the new MAC, while the other constitutes the new MIC (c) (for the sake of simplicity, only one MIC nucleus is shown in the figure; see (66) for more details), while the maternal MAC nucleus is progressively degraded (d). During the development of the new MAC, its chromosomes are amplified by endoreplication, and a fraction of the genome (termed MIC-limited) is eliminated. The distinction between segments that are retained or eliminated relies on a comparison between the maternal MIC and the maternal MAC, via small RNAs produced during meiosis: sequences that are present in the maternal MIC but absent from the maternal MAC are eliminated (c) (14). Hence, even though the two F1 progeny have exactly the same germline genotype (f), their MAC content may differ, depending on the content of their respective maternal MACs (g). The model that we propose relies on the assumption that during interspecific crosses, sequences that are present in the paternal MIC but absent from the maternal MAC are also eliminated. This would explain how crosses between distant species can lead to a viable and fertile offspring. Indeed, let us consider a conjugation event between two genetically divergent species (say A and B), and let us focus on the daughter cell from the A side of the cross (a). Its new MIC is allotetraploid, with the full sets of chromosomes from A and B (b). But given that B sequences are divergent from the A sequences present in its maternal MAC, they will all be eliminated during the development of its new MAC (c). Thus, in the side A daughter cell, the new MAC only expresses genes coming from the A genome, thereby effectively hiding any BDMI with the B genome. Hence, although genetically hybrid (A+B), the side A daughter cell should be viable and retain the phenotype of its maternal lineage (A). Moreover, these cells should be able to complete meiosis normally, as the divergence between the two genomes limits the risk of mispairing between homeologous A/B chromosomes. In F2 individuals resulting from a backcross between an AxB hybrid and an A individual, meiosis is expected to proceed normally for A chromosomes, and hence, even though some of their B chromosomes mis-segregate, these F2’s should lead to a viable and fertile offspring. This model can account for the introgression of DNA coming from B in the MIC-limited compartment of species A.

The third question raised is how the F1 hybrid MIC may have managed to complete meiosis to perpetuate itself. Indeed, the most likely cause of sterility in true F1 hybrids is meiotic failure, due to sequence divergence and/or karyotypic differences between the two parental genomes. We propose that each of these hybridization events was associated with a doubling of ploidy (i.e. with a set of diploid chromosomes from each parent), possibly via endoreplication, as has been described in the related ciliate *Tetrahymena,* after conjugation with a MIC-defective cell (56, 57). It is well established that many whole genome duplications (WGDs) resulted from hybridization events (allopolyploidization) (58–60). When an allopolyploidization involves genetically distant parental genomes, the two sets of duplicated chromosomes can be readily distinguished by the meiotic recombination machinery - thereby allowing proper segregation during meiosis. Allopolyploidizations are thought to be an important source of phenotypic variation (59). But in the present case, the allotetraploid hybrids would retain the original *P. sonneborni* phenotype essentially unchanged, since introgressed genes cannot be expressed. Six independent WGD events have been reported in paramecia (31, 61), which indicates that these events are quite common in that clade. However, if our model is correct, this would imply that WGDs have been exceptionally frequent in the lineage of *P. sonneborni*. One plausible explanation for this high rate is that allopolyploidization mutational events are effectively neutral in *P. sonneborni*, whereas in other taxa, such events are generally counter-selected because of BDMIs between the two parental genomes.

Although our model explains how *P. aurelia* species might escape the deleterious consequences of inter-specific mating, it leaves open further questions. First, it does not provide any clue regarding the *raison d’être* of these recurrent introgressions. Because introgressed genes can hardly be expressed, these events do not appear to contribute much to the adaptive potential of *P. sonneborni*. Second, this model predicts that inter-specific mating can be tolerated only if it involves a species that is genetically distant enough to be recognized as non-self during MAC development and meiosis (Figure 3). Hence, selection should still favor the evolution of traits that prevent inter-specific pairing. A third intriguing point is the fact that this promiscuous behavior is observed only in the *P. sonneborni* lineage. For *P. sonneborni* and *P. sexaurelia*, our experimental design provided the same power to detect HT events originating from subclade A. Yet, whereas we identified ∼6,000 kb of HT segments coming from subclade A in *P. sonneborni,* we detected only 76 kb in *P. sexaurelia* (Fig. 1A,C). Moreover, we detected very few HT segments in the five other MIC genomes analyzed, and they are all short, suggesting that they result from a different mode of horizontal transfer (e.g. via viral vectors) or that they result from much more ancient inter-specific hybridization. So far, *P. sonneborni* appears to be unique among paramecia in showing such a high rate of introgression in its recent ancestry.

We speculate that the large size of the MIC-limited compartment in *P. sonneborni* (135 Mb compared to ∼30 Mb in other *aurelia* species) might result from its propensity to hybridize with distant species. Although the HT segments that we identified in *P. sonneborni* cover only 7.4% of its MIC-limited compartment, they probably represent only the visible tip of the iceberg. Indeed, our study design allowed the detection of HT segments only if they encompassed MAC-destined regions from the donor genome and if they were closely related (>80% sequence identity) to the available reference genomes. Given that sequences in the MIC-limited compartment cannot be expressed, they diverge rapidly, like pseudogenes. It is therefore plausible that a large fraction of the MIC-limited compartment of *P. sonneborni* corresponds to ancient HT segments that are no longer recognizable. Thus, the large size of the genome of *P. sonneborni* might simply reflect the recurrent gains of full sets of chromosomes via allotetraploidization, followed by progressive shrinking through deletions or chromosome loss. Of note, we were hesitant in the use of the term ‘introgression’ to qualify the HT events detected in *P. sonneborni*, because we found no evidence of recombination between homologous loci from the donor and recipient genomes. We eventually chose to use this terminology to highlight the fact that they resulted from sexual processes (mating and formation of a zygote by fusion of gametes). However, if our hypothesis is correct, it might be more accurate to use the term ‘asymmetric allopolyploidization’ to describe such cases of hybridization-mediated WGDs followed by the rapid decay of one of the parental genomes.

The genome of *P. sonneborni* provides an interesting perspective on the definition of species. The model we propose (Figure 3) explains how it is possible for a species to persist, despite recurrent hybridization with very distant lineages. Indeed, owing to nuclear dimorphism, hybrids remain phenotypically identical to their maternal lineage, even if they have received a full set of chromosomes from a very divergent species. Moreover, they also remain interfertile with their parental lineage. Thus, as long as gene flow is maintained in the MAC-destined compartment (which represents only ∼35% of the *P. sonneborni* genome), individuals can be considered as being part of the same species, even if their MIC-limited compartment is not shared. This tentative extension of the definition of a species echoes Sonneborn’s own hesitation in recognizing that what he initially called “varieties” of *P. aurelia* (12) are actually sexually incompatible species (16). Ironically, the promiscuous sibling is the one that bears the name of the father of *Paramecium* genetics.

## Materials and Methods

### Detection of horizontal transfer across aurelia species

We analyzed the MIC genome of each species *(P. tetraurelia, P. octaurelia, P. biaurelia, P. pentaurelia, P. primaurelia, P. sexaurelia* and *P. sonneborni*) (10) to search for segments that have been subject to horizontal transfer (HT) across the *aurelia* complex (see Suppl. Table S4 for a full list of strains and genome assemblies used in this study). For this, the MIC genome of each focal species was cut into 500 bp-long windows, that were compared to all genomes (MAC and MIC, if available) of the 12 other aurelia species with BLASTN (version 2.5.0+; (62)) using the following parameters:

`blastn -query <infile> -db <db> -out <File_Out> -task blastn -evalue 1e-6 - max_target_seqs 1`

For each window, we retained the best blast hit on each of the target genomes. We then selected windows having their best hit with a species that is not a sister-group of the focal species (see phylogenetic tree in Figure 1A). We retained windows with a high level of sequence similarity (>80% identity), significantly higher than with any other species (> mean + 2 standard deviations). These stringent criteria were chosen to identify windows with strong evidence of recent HT.

Then, starting from these ‘seed windows’, we extended the HT segment by aggregating neighbor windows (< 5kb apart) having their best hit on the same scaffold of the non-sister species. The extension was stopped as soon as a seed window matching another scaffold was reached or when the distance to the next window matching the same scaffold exceeded 5 kb. Finally, we retained HT segments that contained at least 4 seed windows, and that included at least 70% of windows having their best hits on the same scaffold. The threshold of 4 seed windows was chosen to ensure that HT segments are at least 2kb-long.

We first performed the HT search by comparing MIC windows of the focal species (query) against the MAC genomes of the 12 other *aurelia* species (targets). We then repeated the analyses using as targets the MIC instead of the MAC genomes, for the 7 species for which both the MAC and the MIC were available. In *P. sonneborni*, this second strategy identified 275 additional candidate HT segments, all matching with the MIC of *P. sexaurelia*. However, given that we only have the MAC but not the MIC genome of *P. jenningsi*, we could not ascertain that these segments were absent from the sister-group. In the other species, both strategies gave very similar results (with at most 8 new HT segments identified). To be conservative, the analyses presented in the main text were restricted to HT segments detected with the first strategy (MAC targets).

To analyze synonymous divergence (dS), we retrieved genome-wide sets of orthologous genes between *P. sonneborni* and other species of the *P. aurelia* complex from the study by Sellis et al. (27). For *P. jenningsi* , which was not included in Sellis’ study, we used the set of 473 one-to-one orthologs reported in another study (17). For each pair of ortholog, we aligned coding sequences based on the protein alignment (build with *Clustal Omega* (*63*) and computed dS with the method of Nei and Gojobori (64) using *SeaView* (*65*). Lists of orthologs and dS values are provided in the Zenodo repository.

### Analysis of HT genes located in the MAC-destined compartment

Based on annotations of the *P. sonneborni* MAC genome assembly (25), we identified 39 gene predictions located in HT segments encompassing MAC-destined regions. For each of them, we used BLASTP to identify homologs in other species of the *P. aurelia* complex. We computed multiple alignments to investigate the conservation of each protein family. In 36 out of 39 cases, we identified nonsense and/or frameshift mutations in the *P. sonneborni* predicted gene with respect to its homologs. We also analyzed the expression level of these predicted genes using RNAseq data from *P. sonneborni* vegetative cell cultures (17).

### Analysis of HT segments in the genome of *P. sonneborni* strain Rio Lg

The *P. sonneborni* strain Rio Lg_Jac 5III was sampled by Sascha Krenek in March 2012, in Lagoa de Jacarepaguá (22° 59’ 7’’ S, 43° 23’ 56’’ W, Rio de Janeiro, Brazil), and kindly sent to us by Alexey Potekhin. In our lab, we checked that this strain belongs to the same species as the *P. sonneborni* reference strain (ATCC 30995): they cross readily, and the F1 hybrids are viable (though we haven’t analyzed the F2 generation). Whole-cell DNA from the Rio Lg strain was extracted from a multi-caryonidal culture obtained by mass autogamy of a single caryonide, after centrifugation and washes with Volvic water. The cell pellet was lysed in 4 volumes of lysis solution (0.5 M EDTA (pH 9); 1% N-lauroylsarcosine; 1% SDS; 1 mg/mL proteinase K) and incubated at 55°C overnight. DNA was then extracted with Tris–HCl-phenol (pH 8) with gentle agitation followed by dialysis against TE (10 mM Tris–HCl; 1 mM EDTA (pH 8)) containing 25% ethanol and then against TE. After RNAse A treatment, followed by a new round of phenol extraction and dialysis against 10 mM Tris (pH 8), an Oxford Nanopore Technology (ONT) library was constructed and sequenced on a P2 solo device using MinKNOW version 25.09.16, sequencing kit SQK-NBD114.24 and an R10.4.1 flowcell. Bases were called with the super accurate mode of dorado 7.11.2. Sequence reads were mapped on the *P. sonneborni* MIC genome assembly (from reference strain ATCC 30995) (10) with minimap2 (version 2.30-r1287):

minimap2 -a -t 40 --MD --cs -c --secondary=no -L -x map-ont psonneborni_mic_ATCC_30995_454Contigs.fa –

We used ‘samtools view’ to exclude from the BAM file all reads showing more than 2% divergence to the reference assembly, and ‘samtools coverage’ to analyze sequencing depth along the genome. We used ‘bedtools coverage’ to count the number of reads overlapping each of the 521 HT segments identified in the ATCC 30995 reference assembly.

## Data availability

All genome and annotation files are available from the NCBI or from the ParameciumDB download section (links and accession numbers are provided in Supplementary Table S4). The complete list of detected HT segments (along with images of their conservation profiles – as in Figure 1B) and R scripts required to reproduce the figures are available on Zenodo (https://doi.org/10.5281/zenodo.18684075). Sequencing data from *P. sonneborni* strain Rio Lg_Jac 5III have been deposited in ENA (accession number ERR17782375).

## Acknowledgments

This work was performed using the computing facilities of the CC LBBE/PRABI. We thank Carina Mugal and Mark Stoneking for helpful comments on the manuscript. We thank Sascha Krenek for providing information on the strain Rio Lg that he had collected, Fransiska Szokoli who initially characterized this strain and Alexey Potekhin who sent it to us. We acknowledge the sequencing and bioinformatics expertise of the I2BC High-throughput sequencing facility, supported by France Génomique (funded by the French National Program "Investissement d’Avenir" ANR-10-INBS-09). This work was supported by the Centre National de la Recherche Scientifique, and by the Agence Nationale de la Recherche (ANR-18-CE12-0005/LaMarque). Work in SD laboratory is supported by the Agence Nationale de la Recherche (ANR-23-CE12-0027; ANR-25-CE12-7757), and by the Fondation de la Recherche Medicale (https://www.frm.org) (Equipe FRM EQU202203014643).

## Author Contributions

FB developed the workflow to detect and classify HT segments. OA and SP were involved in data curation and data analysis. LD, LS and EM performed formal analysis, annotation and data curation. XZ contributed to the sequencing of strain Rio Lg. LD, EM, SD and LS were responsible for conception, supervision and funding of the project. LD wrote the manuscript, with the help of EM, SD and LS. All authors approved the final manuscript.

## Competing Interest Statement

no competing interests.

## Supplementary material

### Supplementary Tables

**Suppl. Table S1:** Number (and percentage) of HT segments covered by sequence reads from strain Rio Lg, in different genomic compartment. The ‘sequencing coverage’ indicates the fraction of each compartment that is covered by at least one read from Rio Lg.

| Reference strain (ATCC 30995) |  | Strain 'Rio Lg' |  |
| --- | --- | --- | --- |
| Compartment | Number of HT segments | Sequencing coverage | Number (%) of HT segments covered |
| MIC-specific (102.6 Mb) | 467 | 6.7% | 42 (9%) |
| MAC-variable (30.4 Mb) | 42 | 50.3% | 13 (31%) |
| MAC-destined (73.9 Mb) | 12 | 93.8% | 10 (83%) |
| Total | 521 |  | 65 (12%) |

**Suppl. Table S2:** Number (and percentage) of HT segments originating from each donor lineage, among the full set of HT segments identified in ATCC 30995, and in the subset retrieved in Rio Lg.

| Reference strain (ATCC 30995) |  | Number (%) of HT segments covered in strain Rio Lg |
| --- | --- | --- |
| Donor lineage | Number (%) of HT segments |  |
| *P. novaurelia* | 17 (3%) | 0 (0.0%) |
| *P. quadecaurelia* | 117 (22%) | 20 (31%) |
| *P. tredecaurelia* | 169 (32%) | 26 (40%) |
| *P. sexaurelia* | 218 (42%) | 19 (29%) |

**Suppl. Table S3:**
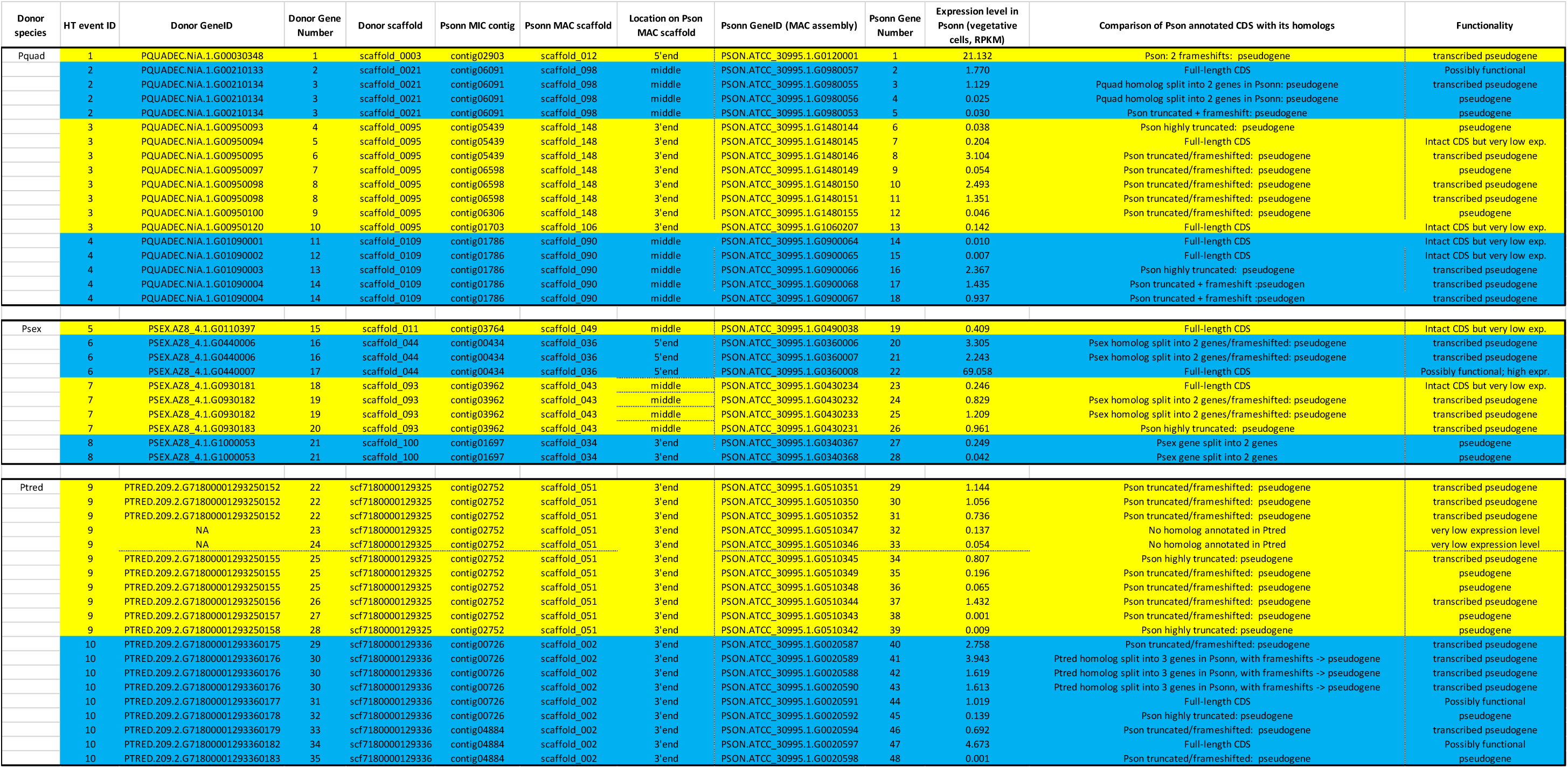
List of genes identified in HT segments located in the MAC-destined compartment of *P. sonneborni*.

**Suppl. Table S4:** List of genome assemblies and strains used in this study.

| Species | Species code | Assembly name | Category | Strain | NCBI Assembly Accession/ParameciumDB link |
| --- | --- | --- | --- | --- | --- |
| *P. biaurelia* | *pbia* | pbiaurelia_mac_V1-4_v1.0 | MAC | V1-4 | https://paramecium.i2bc.paris-saclay.fr/download/Paramecium/biaurelia/V1-4/sequences/ |
| *P. biaurelia* | *pbia* | pbiaurelia_mic_V1-4_v1.0 | MIC | V1-4 | GCA_980646935.1 |
| *P. decaurelia* | *pdec* | pdecaurelia_mac_223_v1.0 | MAC | 223 | https://paramecium.i2bc.paris-saclay.fr/download/Paramecium/decaurelia/223/sequences/ |
| *P. dodecaurelia* | *pdod* | pdodecaurelia_mac_274_v1.0 | MAC | 274 | https://paramecium.i2bc.paris-saclay.fr/download/Paramecium/dodecaurelia/274/sequences/ |
| *P. jenningsi* | *pjen* | pjenningsi_mac_M_v1.0 | MAC | M | https://paramecium.i2bc.paris-saclay.fr/download/Paramecium/jenningsi/M/sequences/ |
| *P. novaurelia* | *pnov* | pnovaurelia_mac_TE_v1.0 | MAC | TE | https://paramecium.i2bc.paris-saclay.fr/download/Paramecium/novaurelia/TE/sequences/ |
| *P. octaurelia* | *poct* | poctaurelia_mac_138_v1.0 | MAC | 138 | https://paramecium.i2bc.paris-saclay.fr/download/Paramecium/octaurelia/138/sequences/ |
| *P. octaurelia* | *poct* | poctaurelia_mic_138_v1.1 | MIC | 138 | GCA_980644265.1 |
| *P. pentaurelia* | *ppen* | ppentaurelia_mac_87_v1.0.MAC-constitutive | MAC | 87 | https://paramecium.i2bc.paris-saclay.fr/download/Paramecium/pentaurelia/87/sequences/ |
| *P. pentaurelia* | *ppen* | ppentaurelia_mic_87_v1.1 | MIC | 87 | GCA_980644265.1 |
| *P. primaurelia* | *ppri* | pprimaurelia_mac_AZ9-3_v1.1.MAC-constitutive | MAC | AZ9-3 | https://paramecium.i2bc.paris-saclay.fr/download/Paramecium/primaurelia/AZ9-3/sequences/ |
| *P. primaurelia* | *ppri* | pprimaurelia_mic_AZ9-3_v1.0 | MIC | AZ9-3 | GCA_980642815.1 |
| *P. quadecaurelia* | *pqua* | pquadecaurelia_mac_NiA_v1.0 | MAC | NiA | https://paramecium.i2bc.paris-saclay.fr/download/Paramecium/quadecaurelia/NiA/sequences/ |
| *P. sexaurelia* | *psex* | psexaurelia_mac_AZ8-4_v1.1 | MAC | AZ8-4 | https://paramecium.i2bc.paris-saclay.fr/download/Paramecium/sexaurelia/AZ8-4/sequences/ |
| *P. sexaurelia* | *psex* | psexaurelia_mic_AZ8-4_v1.1 | MIC | AZ8-4 | GCA_980645855.1 |
| *P. sonneborni* | *pson* | psonneborni_mac_ATCC_30995_v1.0.MAC-constitutive | MAC | ATCC_30995 | https://paramecium.i2bc.paris-saclay.fr/download/Paramecium/sonneborni/ATCC_30995/sequences/ |
| *P. sonneborni* | *pson* | psonneborni_mic_ATCC_30995_454Contigs | MIC | ATCC_30995 | GCA_980648905.1 |
| *P. tetraurelia* | *ptet* | ptetraurelia_mac_51_v2.0.MAC-constitutive | MAC | 51 | https://paramecium.i2bc.paris-saclay.fr/download/Paramecium/tetraurelia/51/sequences/ |
| *P. tetraurelia* | *ptet* | ptetraurelia_mic_51_v1.2 | MIC | 51 | https://paramecium.i2bc.paris-saclay.fr/download/Paramecium/tetraurelia/51/sequences/ |
| *P. tredecaurelia* | *ptre* | ptredecaurelia_209_AP38_filtered | MAC | 209 | https://paramecium.i2bc.paris-saclay.fr/download/Paramecium/tredecaurelia/209_AP38/sequences/ |

### Supplementary Figures

**Suppl. Figure S1:**
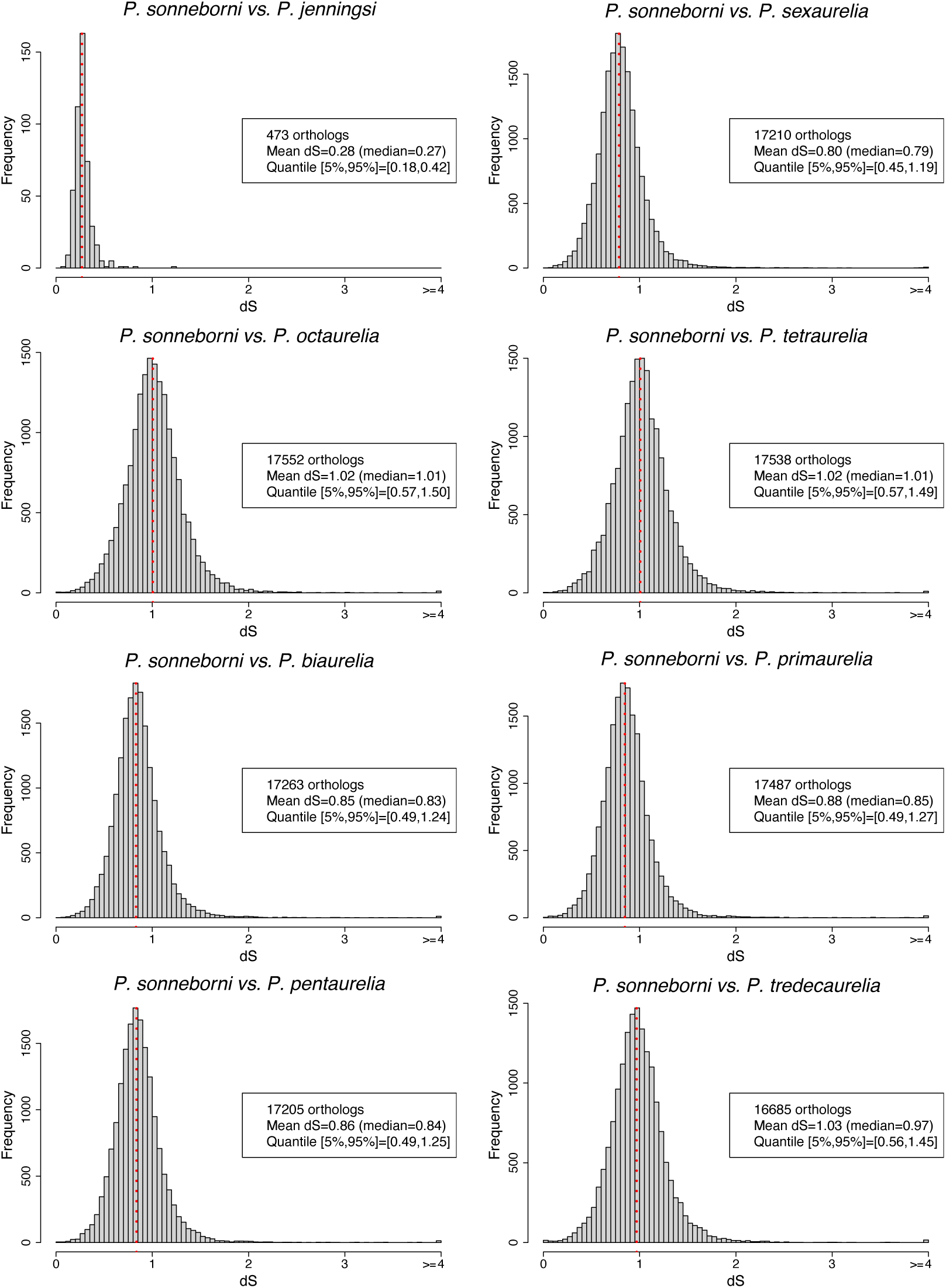
Distribution of synonymous divergence between *P. sonneborni* and other species of the *P. aurelia* complex. For each species pair, the synonymous divergence (dS) was computed on genome-wide sets of orthologous genes from Sellis et al. (*PLOS Biol.* **19**, e3001309 2021), except for *P. jenningsi*, which was not present in their study, and for which we used the set of 473 single-copy orthologs collected by Sawka-Gadek et al. (*Genome Biol. Evol.* **13** 2021).

**Suppl. Figure S2:**
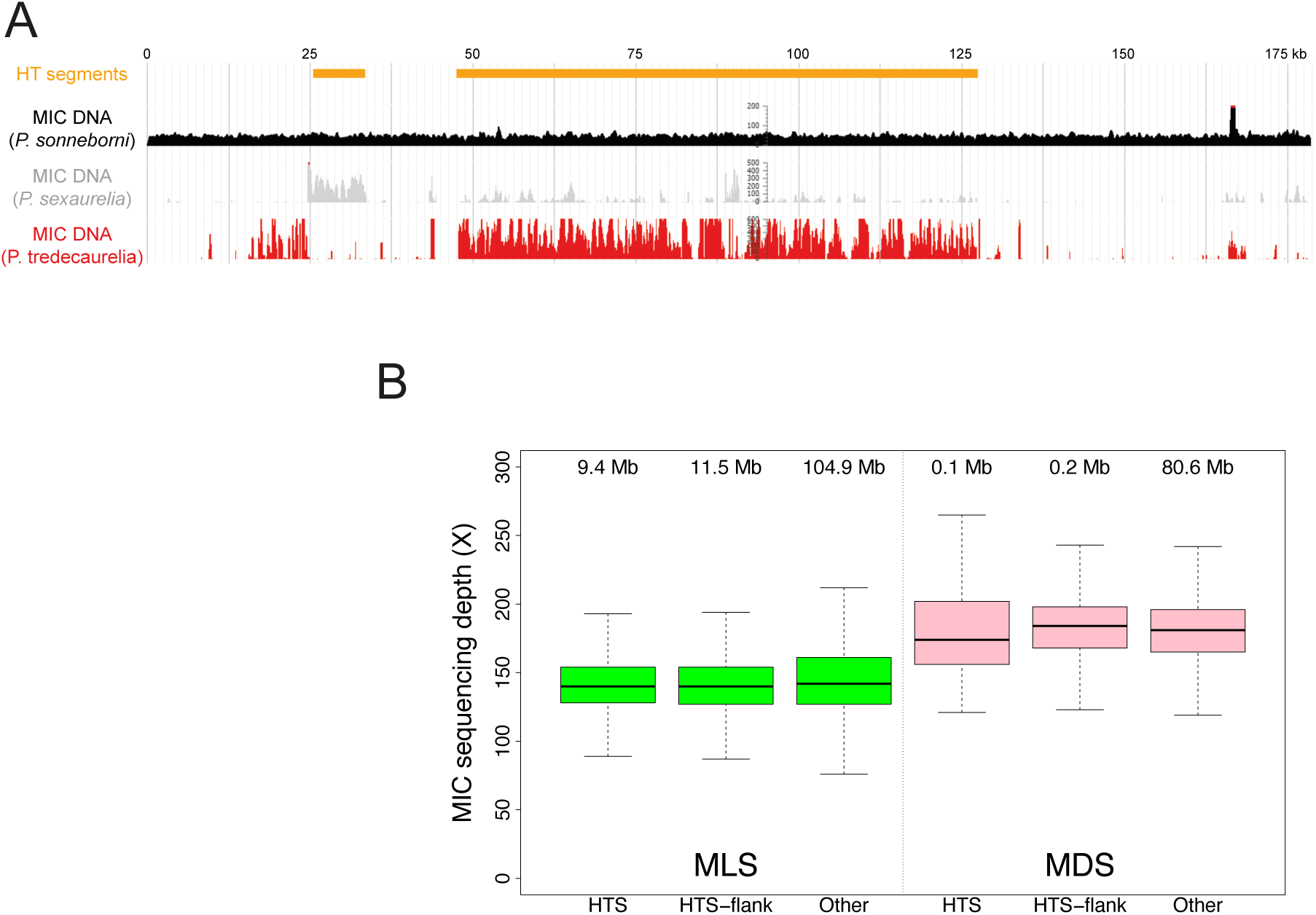
Evidence that HT segments detected in *P. sonneborni* do not result from DNA contamination artefacts. **A.** Example of a MIC contig of *P. sonneborni* (contig00015) that includes two HT segments (orange rectangles), the first derived from *P. sexaurelia* (scaffold_001.2), the second from *P. tredecaurelia* (scf7180000129386). If HT segments resulted from DNA contamination, this would imply that contig00015 corresponds to an artefactual chimeric assembly of DNA fragments coming from three sources (*P. sonneborni*, *P. sexaurelia* and *P. tredecaurelia*). Hence, the sequencing depths would be expected to differ between the two HT segments (originating from *P. sexaurelia* and *P. tredecaurelia*), and to be lower in HT segments than in flanking regions (*P. sonneborni*). The first profile (black) indicates the sequencing depth for the *P. sonneborni* MIC DNA sample. This profile shows a uniform sequencing depth along the whole contig, including in regions outside of HT segments, in contradiction with the hypothesis of a DNA contamination artefact. As a control, we also mapped sequence reads from MIC DNA samples obtained from *P. sexaurelia* (grey) or *P. tredecaurelia* (red). As expected, we observed a high density of reads from each species mapping on the corresponding HT segment. The heterogeneity of coverage reflects the fact that the sequences of the *P. sexaurelia* and *P. tredecaurelia* strains that we used in this study are not identical to the HT segments present in *P. sonneborni*, again in contradiction with the hypothesis that these HT segments might result from DNA contamination. **B.** Distribution of sequencing depth within and outside HT segments in *P. sonneborni*. The MIC genome assembly was split into 500bp-long non-overlapping windows, which were classified as MIC-limited (MLS) or MAC-destined (MDS), based on their MAC sequencing depth, and then subdivided into three subsets: windows that overlap a HT-segment (HTS), windows that flank a HT-segment (HTS-flank), and windows located on other MIC contigs (i.e. not flanking a HT-segment). Both in MLS and in MDS regions, the distribution MIC sequencing depth (represented by boxplots) is the same within HT-segments as in their flanking regions and the same as in the rest of the genome, consistent with the assumption that HT-segments correspond to authentic *P. sonneborni* MIC DNA. The amount of DNA in each class of windows is given above the corresponding boxplot.

**Suppl. Figure S3:**
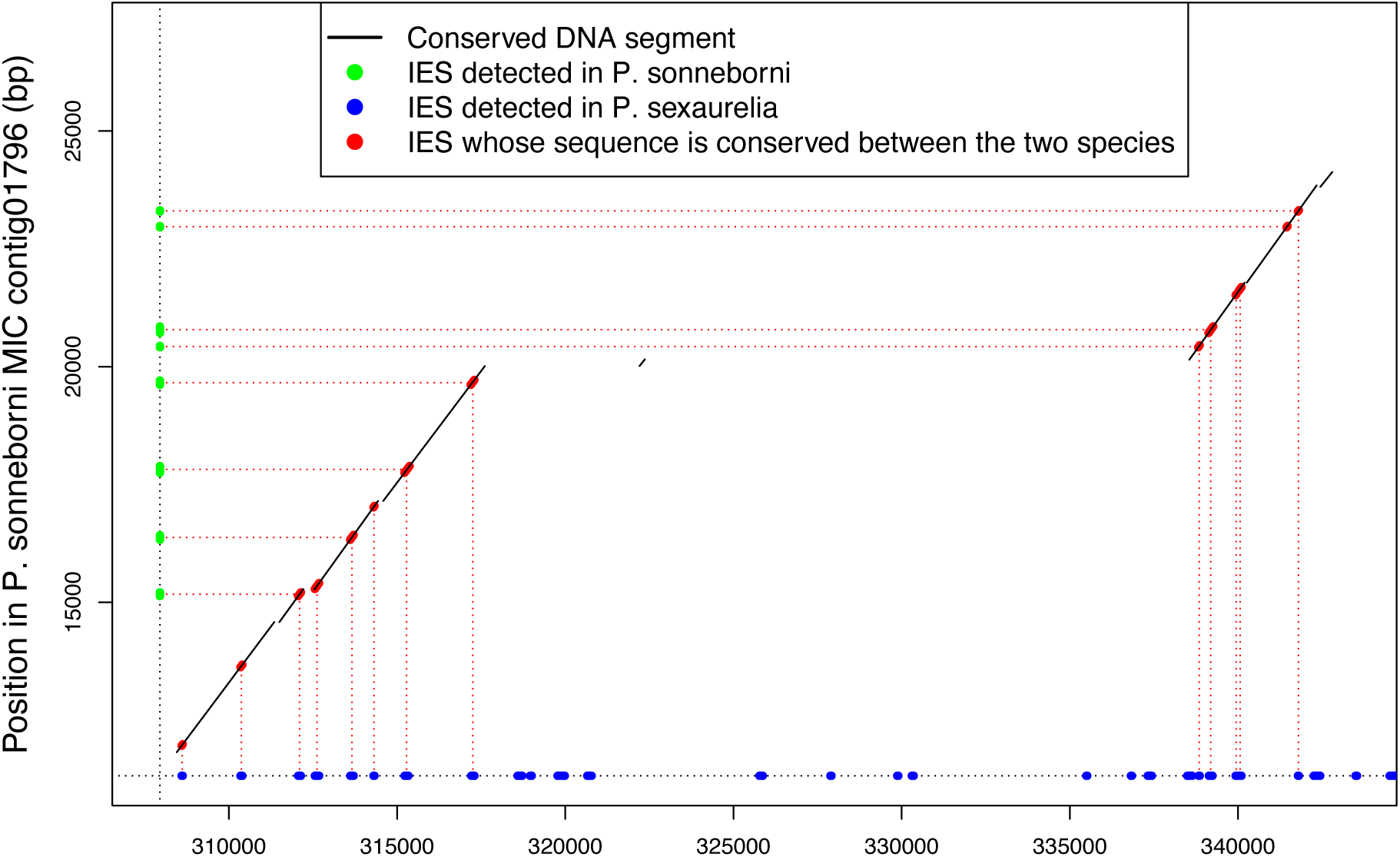
Presence of conserved IESs within HT segments. This figure presents a dot plot alignment between a *P. sonneborni* MIC contig (contig01796) and the *P. sexaurelia* MIC scaffold (scaffold_103), with which we identified a HT segment (88% sequence identity between the two species). IESs have been previously detected in the MIC genomes of *P. sonneborni* and of *P. sexaurelia* (24). The figure shows that this HT segment encompasses 14 conserved IESs, among which 7 were detected in both species, 6 were detected only in *P. sexaurelia* and 1 only in *P. sonneborni*. Given that IESs are absent from the MAC genome, the presence of conserved IESs within the HT segment indicates that it resulted from the transfer of MIC, rather than MAC DNA.

## Notes

https://doi.org/10.5281/zenodo.18684075

### Competing Interest Statement

The authors have declared no competing interest.

### Summary of Updates

We corrected an accession number in the data availability section. We slightly modified the introduction and the discussion.

